# Autocatalytic microtubule nucleation determines the size and mass of spindles

**DOI:** 10.1101/174078

**Authors:** Franziska Decker, David Oriola, Benjamin Dalton, Jan Brugués

## Abstract

Regulation of size and growth is a fundamental problem in biology. A prominent example is the formation of the mitotic spindle, where protein concentration gradients around chromosomes are thought to regulate spindle growth by controlling microtubule nucleation (*1*, *2*). Previous evidence suggests that microtubules nucleate throughout the spindle structure (*3*-*5*). However, the mechanisms underlying microtubule nucleation and its spatial regulation are still unclear. Here, we developed an assay based on laser ablation to directly probe microtubule nucleation events in *Xenopus laevis* egg extracts. Combining this method with theory and quantitative microscopy, we show that the size of a spindle is controlled by autocatalytic growth of microtubules, driven by microtubule-stimulated microtubule nucleation. The autocatalytic activity of this nucleation system is spatially regulated by the limiting amounts of active microtubule nucleators, which decrease with distance from the chromosomes. Thus, the size of spindles is determined by the distance where one microtubule nucleates on average less than one new microtubule. This mechanism provides an upper limit to spindle size even when resources are not limiting and may have implications for spindle scaling during development (*6*, *7*).

## Main text

A general class of problems in biology is related to the emergence of size and shape in cells and tissues. Reaction diffusion mechanisms have been broadly successful in explaining spatial patterns in developmental biology as well as some instances of intracellular structures (*8*, *9*). The mitotic spindle, a macromolecular machine responsible for segregating chromosomes during cell division, is thought to be a classic example of such reaction diffusion processes. A diffusible gradient of the small GTPase Ran emanating from chromosomes has been shown to trigger a cascade of events that result in the nucleation of microtubules, the main building blocks of the spindle (*1*, *2*). The spatial distribution of microtubule nucleation is key for understanding size and architecture of large spindles. This is because microtubules in these spindles are short and turnover rapidly (*3*, *10*, *11*). The mechanisms underlying the spatial regulation of microtubule nucleation, however, are still unclear (*12*, *13*). One possibility is that the interplay between Ran-mediated nucleation and microtubule turnover governs spindle assembly (*1*, *2*). However, the role of the Ran gradient in determining spindle size is still controversial. For instance, in cell culture systems, the length scale of the Ran gradient does not correlate with spindle size (*5*). A second possibility is that autocatalytic growth accounts for spindle assembly via microtubule-stimulated microtubule nucleation (*4*, *14*-*16*). However, autocatalytic mechanisms suffer from the fact that their growth is hard to control. Although autocatalytic growth can be regulated by limiting the catalyst, such mechanisms are unlikely to function in the large cells of developing eggs such as *Xenopus*, where resources are not limiting (*17*). Understanding the role of microtubule nucleation in setting the size of spindles is limited by the fact that little is known about the rate, distribution, and regulation of microtubule nucleation in spindles (*12*, *13*). This is partly because of the lack of methods to measure microtubule nucleation in spindles.

Microtubules grow from the plus ends while minus ends remain stable (*18*). Thus, the location of minus ends functions as a marker for microtubule nucleation. However, in spindles microtubules constantly flux towards the poles (*19*), and measuring the location of a microtubule minus end at a particular time does not correspond to its original site of nucleation (*3*). To decouple microtubule transport from microtubule nucleation, we inhibited kinesin-5 (Eg5) in spindles assembled in *Xenopus laevis* egg extracts. This inhibition stops microtubule transport and leads to the formation of radially symmetric monopolar spindles (monopoles) that have a similar size as regular spindles (*20*, *21*) (Fig. 1A and S1). The location of minus ends in these monopoles exactly corresponds to the location of microtubule nucleation.

**Fig. 1:**
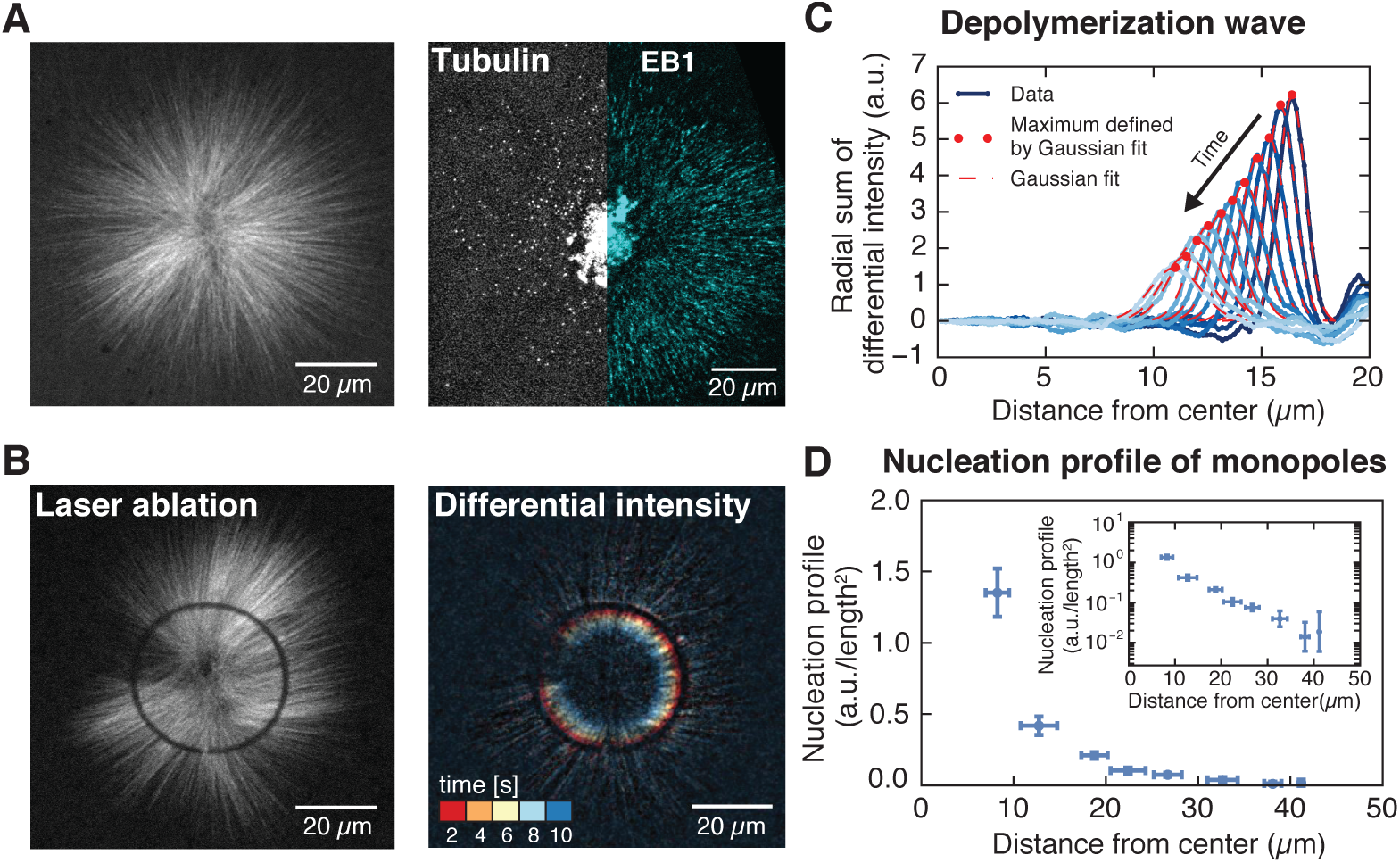
Microtubule nucleation in monopolar spindles is spatially regulated. (A) Fluorescence image of a monopolar spindle (left), and single-molecule fluorescent tubulin and EB1-GFP (right). (B) Circular laser cut and corresponding differential intensity depolymerization front at different times. (C) Radial sum of differential intensities at different time points (from dark to light blue) of one cut at a radius of 19 μm from the center. The area under each curve equals the mass of microtubules depolymerized per time interval of 2 s. (D) Nucleation profile of monopolar spindles (N = 117 cuts, mean ± SD).

Three independent measurements show that inhibiting microtubule transport does not affect dynamic parameters of microtubules. First, microtubules in these structures polymerize at 20.9 ± 5.1 *μ*m/min (N = 7 monopoles, Fig. 1A and movie S1), which is indistinguishable from the polymerization velocity in spindles, 22.7 ± 8.4 *μ*m/min (N = 4 spindles). Second, microtubules from monopolar and control spindles depolymerize at the same velocity (33.5 ± 6.4 *μ*m/min and 35.9 ± 7.3 *μ*m/min respectively, see Fig. S2). Third, microtubule lifetime distributions of monopolar spindles, measured by single-molecule microscopy of tubulin dimers, give an average lifetime of 19.8 ± 2.2 s, consistent with similar measurements in regular spindles (*11*) (materials and methods and movie S2).

To localize microtubule nucleation events, we measured the density of minus ends in monopolar spindles by analyzing synchronous waves of microtubule depolymerization from laser cuts similar to (*3*). Briefly, the minus end density at the location of the cut can be obtained from the decrease of the microtubule depolymerization wave, but as opposed to Ref. (*3*), our method resolves the minus end locations with a single laser cut, see Fig. 1B-C, Fig. S3, movie S3 and S4, and materials and methods. We measured the microtubule nucleation profile across the entire structure by performing laser cuts at different distances from the center of the monopoles. These measurements revealed that microtubule nucleation extends throughout monopoles, with highest nucleation near the center and monotonically decreasing far from the center (see Fig. 1D), indicating that the strength of microtubule nucleation is spatially regulated.

Several mechanisms have been proposed to regulate microtubule nucleation. From a biophysical perspective, these mechanisms can be categorized into two scenarios: (1) microtubule-dependent nucleation, in which a pre-existing microtubule stimulates the nucleation of a new microtubule, or (2) microtubule-independent nucleation, in which factors other than pre-existing microtubules (e.g. diffusible cues in the cytoplasm) stimulate nucleation (*12*-*15*, *22*-*24*).

If microtubule nucleation depends on pre-existing microtubules, altering microtubule stability should change the nucleation profile—a microtubule that exists for a longer time would have a higher probability to stimulate the creation of more microtubules. To test this scenario, we increased microtubule stability by inhibiting the depolymerizing kinesin MCAK (*25*) using antibodies. MCAK inhibition led to a dramatic increase in monopole size (see Fig. 2A). Both the average length and stability of microtubules increased threefold after inhibition (Fig. 2B and C) as assessed by laser ablation (8.0 ± 0.3 *μ*m versus 23.6 ± 3.6 *μ*m, see materials and methods and (*3*)) and single microtubule lifetime imaging (19.8 ± 2.2 s versus 60.4 ± 4.4 s), movie S2 and S5. These measurements are consistent with MCAK modifying the catastrophe rate (*25*, *26*). We measured microtubule nucleation in this perturbed condition and found that the nucleation profile extends further from the center of the monopole, has a larger amplitude, and decays over a larger distance with respect to control monopoles (Fig. 2D). Therefore, the number and spatial distribution of nucleated microtubules does indeed scale with microtubule stability in monopolar spindles, which is inconsistent with microtubule-independent nucleation. Thus, microtubule nucleation in these structures depends on the presence and dynamics of microtubules.

**Fig. 2:**
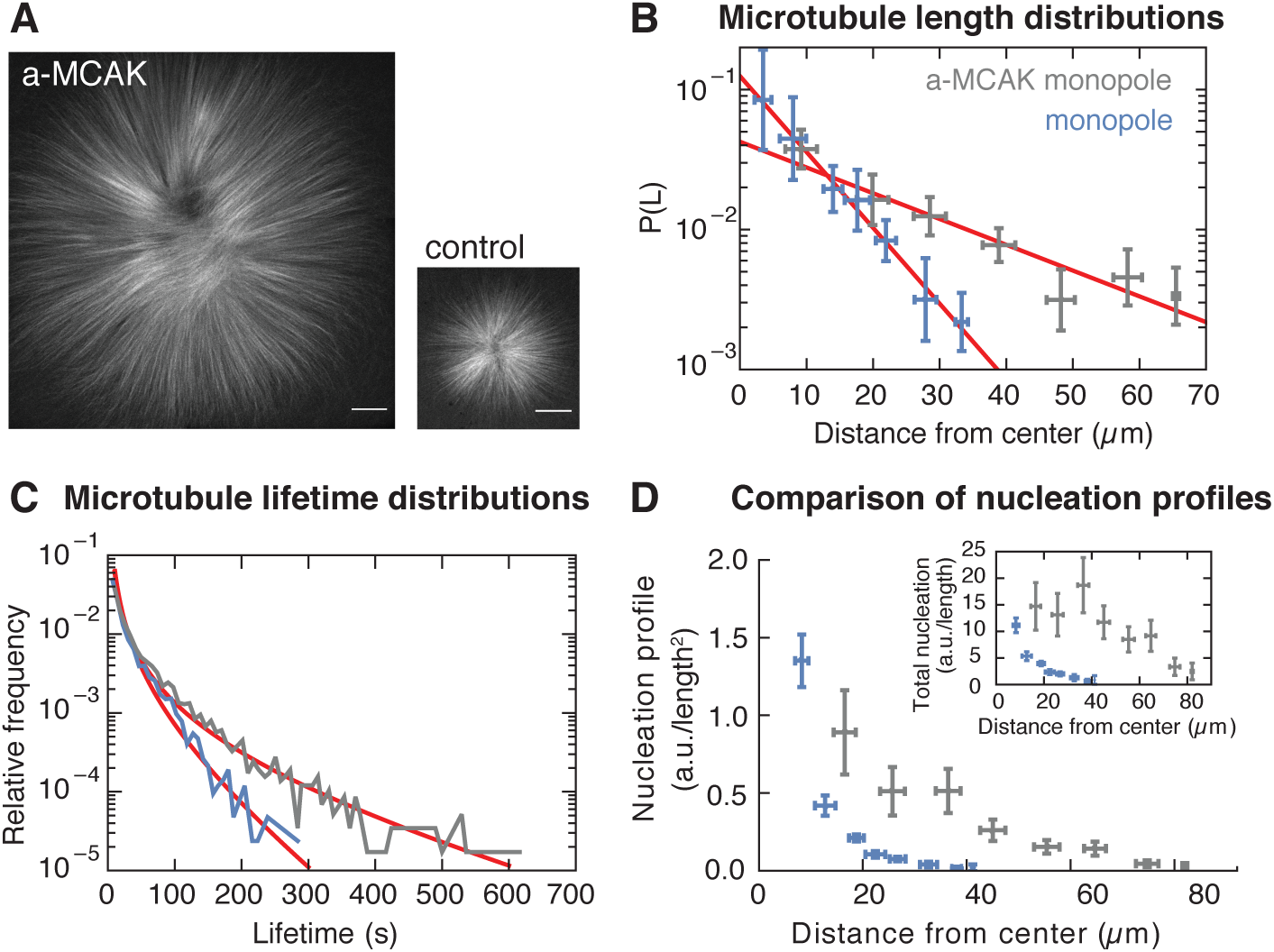
Microtubule nucleation depends on the stability of microtubules. (A) Inhibition of MCAK leads to larger steady-state monopoles. Scale bar = 20 *μ*m (B) Microtubule length distributions measured from laser ablation and fitted to an exponential (mean ± SD). (C) Normalized histograms of microtubule lifetimes of control (N = 5331 speckles, 5 structures) and MCAK-inhibited monopoles (N = 7289 speckles, 3 structures), and corresponding first-passage time fits (see supplementary methods). (D) Nucleation profile of control (N = 117 cuts) and MCAK-inhibited monopoles (N = 74 cuts) (mean ± SD). The inset shows both nucleation profiles multiplied by the circumference length at each radius, which corresponds to the total microtubule nucleation at that distance from the center of the monopole.

The presence and dynamics of microtubules could alter microtubule nucleation in two ways: microtubules could nucleate indiscriminately in the cytoplasm without requiring microtubules, but their presence concentrate active nucleators through transient interactions with microtubules as previously proposed (*5*), or alternatively, microtubules could directly nucleate new microtubules, requiring active nucleators to bind to microtubules to initiate nucleation. In the latter case, the presence of a microtubule is essential for the nucleation process, whereas in the former, microtubules can still nucleate in the absence of microtubules. To test whether microtubule nucleation requires physical proximity to pre-existing microtubules (e.g., a branching process (*14*)), we locally blocked microtubule polymerization by adding inert obstacles near the center of monopoles, at locations where nucleation should be expected according to our measurements (Fig. 2D and E). These localized obstacles cannot prevent the diffusion of nucleators, but would prevent microtubules that polymerize towards them to extend further. Consistent with microtubule-stimulated nucleation, the presence of these obstacles completely inhibited nucleation of new microtubules behind the obstacles, as in a shadow cast by light, whereas microtubules nucleated further around the obstacles, creating a sharp boundary see Fig. 3A. These results suggest that monopolar spindles grow to a size larger than an individual microtubule by microtubule-stimulated microtubule nucleation in physical proximity to pre-existing microtubules, which creates an autocatalytic wave of microtubule growth.

**Fig. 3:**
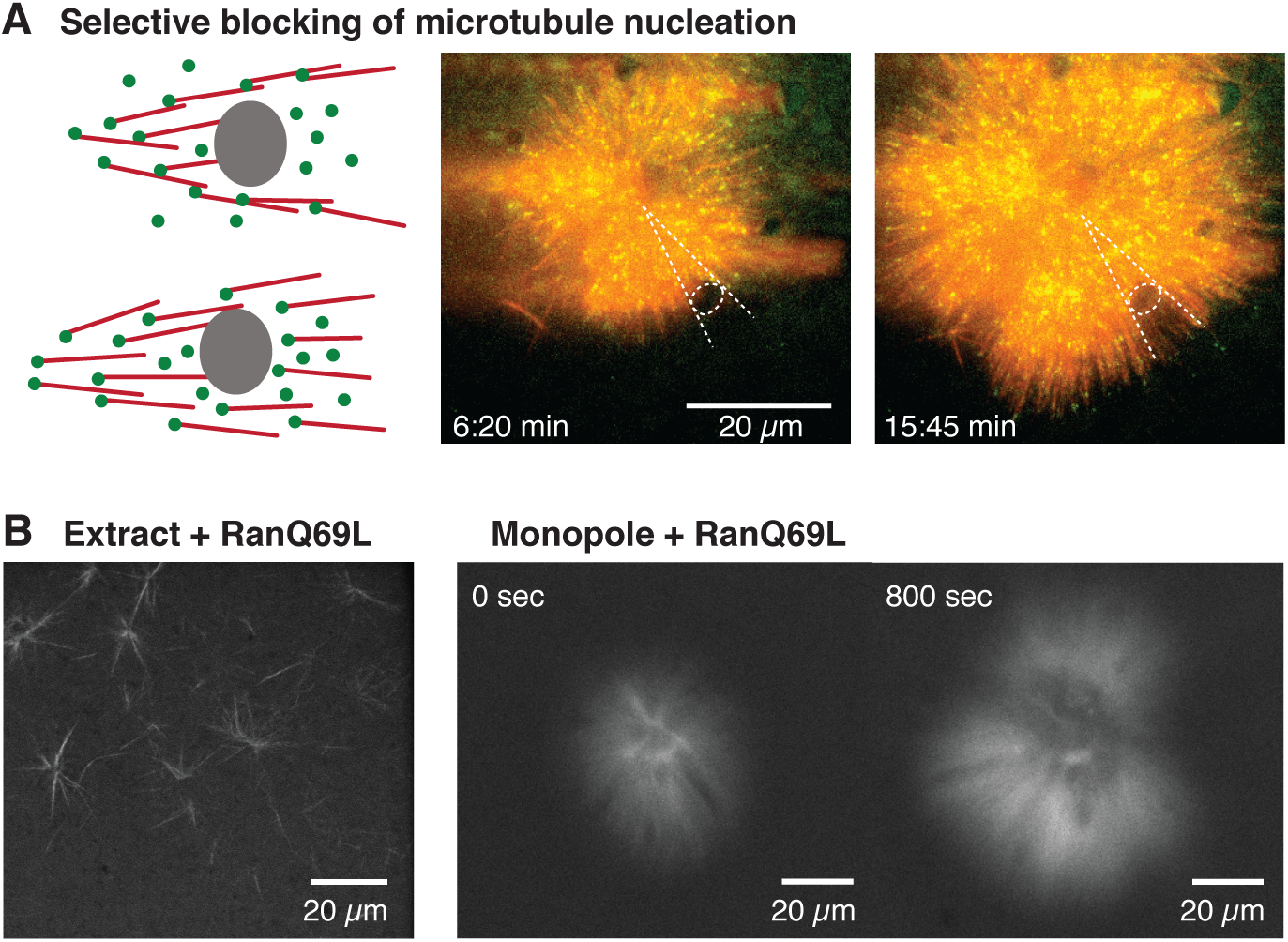
Microtubule nucleation requires phyiscal proximity to pre-exisiting microtubules. (A) Inert fluorocarbon oil microdroplets immobilized to coverslips selectively block microtubule nucleation. Left: Schematic outcomes depending on whether new microtubules are nucleated in phyiscal proximity to pre-existing microtubules (top) or not (bottom). Middle, right: Two time points of a microtubule structure with labeled tubulin (red) and EB1-GFP (green) growing around immobilized frog sperm. The oil microdroplet is highlighted with a dashed ellipse. (B) Left: Addition of RanQ69L to extract homogeneously activates nucleation and creates mini asters after ∼ 20 min (22). Middle, right: Addition of RanQ69L to monopoles leads to immediate growth of new microtubules from the pre-existing monopoles.

For a microtubule structure to have a finite size through an autocatalytic process, each microtubule at the periphery must create on average less than one microtubule at steady state, otherwise the number of microtubules would increase exponentially and the structure would grow unbounded (*16*). However, measurements of the temporal evolution of microtubule mass in spindles show indeed an initial phase of exponential growth (Fig. S4 and (*22*, *27*)). This is also consistent with the observation of microtubules creating more than one microtubule on average when inducing bulk microtubule branching by adding TPX2 and constitutively active Ran (RanQ69L) in extracts (*14*). These observations raise the question of how spindles reach a finite size through autocatalytic growth (as in the control and MCAK-inhibited monopoles). One possibility is that microtubule dynamics change as a result of limiting amounts of tubulin or microtubule-associated proteins (*6*, *7*). However, since our cell-free system is not confined, availability of tubulin and microtubule-associated proteins is not limiting. Furthermore, inhibiting MCAK leads to microtubule growth with a polymerization velocity that is indistinguishable from control monopoles (20.9 ± 5.1 *μ*m/min and 18.8 ± 5.4 *μ*m/min respectively, movie S1 and S6), suggesting that the availability of tubulin appears not to be diffusion-limited. Finally, microtubule dynamics do not change spatially throughout MCAK-inhibited monopoles (Fig. S5), indicating that spatial variations of tubulin amount or microtubule dynamics cannot explain the finite size of these structures.

Another possibility is that microtubule nucleation is limiting. It has been shown that the assembly of spindles requires RanGTP. RanGTP is created only in the vicinity of chromosomes (through the ran nucleotide exchange factor RCC1) which in turn releases spindle assembly factors (SAFs) responsible for nucleating microtubules (*1*, *2*). Since the active SAFs are naturally limited by their spatially restricted generation, a limiting amount of an active microtubule nucleation factor would therefore be a good candidate as the limiting component for both autocatalytic growth and size regulation. To test this idea, we added constitutively active Ran (RanQ69L), to pre-existing monopolar spindles. A limiting pool of active nucleators implies that (i) activating nucleators everywhere in the cytoplasm would lead to unbounded microtubule growth in the monopole (similar to large interphase asters in embryos (*28*)), and (ii) new microtubules should nucleate from the pre-existing microtubules of the structure. Adding RanQ69L to pre-existing monopoles immediately started nucleation of new microtubules preferentially at the edge of the pre-existing structures in a wave-like fashion, consistent with microtubule-stimulated growth (Fig. 3B and movie S7 and S8). This result further suggests that other limiting components that regulate microtubule dynamics alone cannot account for this growth. Taken together, these measurements show that the amount of active nucleators, which is limited by the availability of RanGTP, limits the size of monopolar spindles and is responsible for the bounded growth of these structures.

To test whether a limited pool of active nucleators can quantitatively account for the size and microtubule nucleation in these microtubule structures, we developed a biophysical model of autocatalytic microtubule nucleation (see Fig. 4A and materials and methods). In our model, inactive nucleators are present throughout the cytoplasm and can be activated at the surface of chromosomes, which is a simplification of the activation of SAFs by RanGTP. The total amount of active nucleators depends on the balance between the rate of activation at the chromosomes and the rate of inactivation (accounting for sequestration, hydrolysis, or other processes). Once activated at the chromosomes, nucleators can diffuse in the cytoplasm, bind, and unbind from microtubules. When bound to microtubules, active nucleators can nucleate new microtubules at a certain rate, and the newly nucleated microtubules maintain the same polarity as the mother microtubule (*14*). This process leads to an autocatalytic wave as a consequence of the self-replicating activity of an extended object where, in contrast to a reaction diffusion process, front propagation is independent of microtubule diffusion and it only depends on microtubule dynamics.

**Fig. 4:**
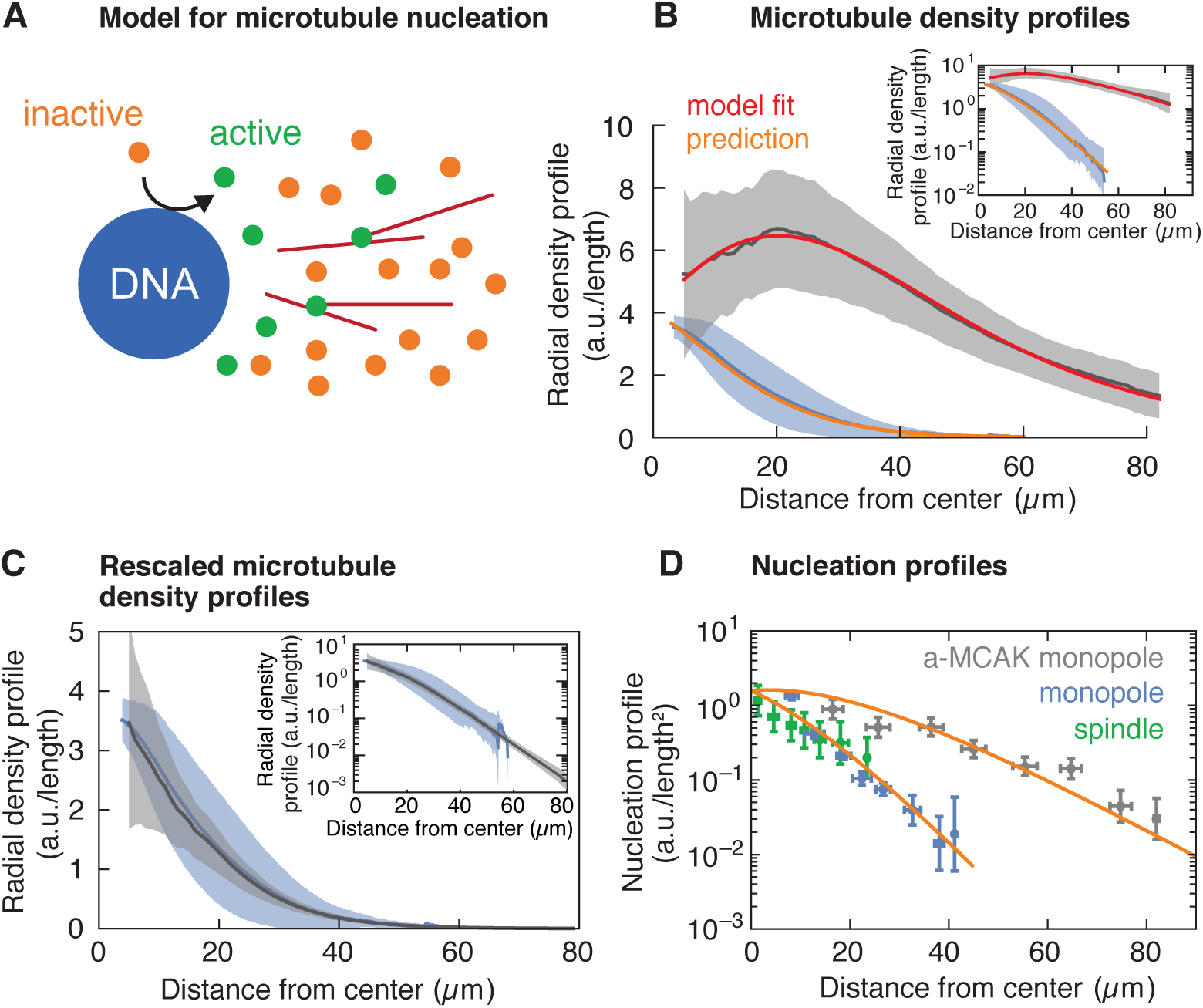
Model for microtubule-stimulated nucleation. (A) Inactive nucleators (orange circles) are activated around DNA. As active nucleators diffuse (green circles), they can bind and unbind microtubules (red lines). Once bound, they can nucleate a new microtubule with a certain probability. Active nucleators become inactive at a constant rate. (B) Radial microtubule density profiles measured from fluorescent images (N_monopoles_ = 40 (blue), N_a-MCAK_ = 18 (gray)) and corresponding model fit to the MCAK-inhibited and prediction to control monopoles. (C) Parameter-free rescaling of the microtubule density profiles: ρ_C_ = ρ_M_ exp[(1/l_M_ -1/l_C_)x], where ρ_C_, l_C_ and ρ_M_, l_M_ are the density and length of microtubules for the control (blue) and MCAK-inhibited (gray) structures, respectively, and x is the distance from the center of the structure. (D) Data and predictions (orange) for the nucleation profiles of control (blue) and MCAK monopoles (gray) up to a global nucleation amplitude, and flux-corrected regular spindles (green) (mean ± SD, N_control_ = 117, N_a-MCAK_ = 74, N_spindle_ = 36 cuts).

In our model, the amount and dynamics of active nucleators should be the same for both control and MCAK-inhibited monopoles, leading to the prediction that the two microtubule density profiles would only differ in a parameter controlling the microtubule length or lifetime. In particular, both profiles should scale to each other without any fitting parameters by changing the microtubule length as measured independently by laser ablation. To test this prediction, we measured the radial profile of microtubule density of control and MCAK-inhibited monopoles (Fig. 4B): These microtubule density profiles are qualitatively different—the density of MCAK-inhibited monopoles increases initially and decreases after reaching a maximum, whereas the control monopole decreases monotonically from the origin. Remarkably, both profiles collapse into each other after the parameter-free rescaling of the MCAK-inhibited monopole (Fig. 4C). To test the model beyond scaling, we fit the MCAK-inhibited profile with the other two remaining parameters and the arbitrary amplitude of the microtubule density profile, which agrees quantitatively with the data (Fig. 4B). By fixing all parameters to the values obtained by this fit and using the measured average microtubule length, the model predicts the control monopole microtubule profile. Finally, we can also predict the MCAK-inhibited and control microtubule nucleation profiles from the fitted parameters up to an arbitrary amplitude (common for both profiles) (Fig. 4D). Remarkably, this prediction is also consistent with flux-corrected microtubule nucleation in regular spindles obtained by laser ablation (see materials and methods, Fig. 4D green circles, movie S9 and S10), showing that the same nucleation mechanism holds for regular spindles. Thus, our model for autocatalytic microtubule nucleation accounts for both the microtubule density and nucleation profiles.

Our data and model are consistent with an autocatalytic mechanism in which microtubule-stimulated microtubule nucleation controls spindle growth. This process is spatially regulated by a gradient of active nucleators that is established by the interplay between the Ran gradient and microtubule dynamics. Microtubules regulate the nucleator activity because they act as the substrate where active nucleators need to bind to nucleate microtubules. Chromatin acts as a trigger for an autocatalytic wave of microtubule nucleation, and at the same time limits spindle size by controlling the amount of active nucleators through RanGTP. This suggests that the amount of active Ran can tune spindle length, and resolves its controversial relation to spindle length regulation: while a diffusion and inactivation process has a characteristic length scale independent of the amplitude of the gradient—set by the ratio of the squared root of the diffusion and inactivation rate—here we show that both the length scale and amplitude of the gradient of nucleators are involved in regulating the size and mass of spindles. Since the length scale of the gradient is amplified by microtubule-stimulated nucleation, the relevant length scale for setting the size is the distance at which a microtubule generates one or fewer microtubules. Our proposed mechanism therefore allows regulation of spindle size and mass by two means, although microtubule nucleation is the principal control parameter, microtubule dynamics can still fine tune the spindle length (*29*).

An autocatalytic nucleation process implies that microtubule structures are capable of richer dynamical behaviors than those arising from the classic view of random nucleation in the cytoplasm via a diffusible gradient. Beyond producing finite-sized structures like spindles and ensuring that new microtubules keep the same polarity as the pre-existing ones, it also allows for a rapid switch into unbounded wave-like growth if nucleators become active throughout the cytoplasm. Indeed, the growth of large interphase asters has been hypothesized as a chemical wave upon Cdk1 activation (*30*, *31*). These properties, characteristic of excitable media, provide a unified view of the formation of spindles and large interphase asters in embryos (*23*) within a common nucleation mechanism. However, microtubule nucleation differs from regular autocatalytic processes in reaction-diffusion systems such as Fisher-waves and Turing mechanisms (*8*, *32*) in that its growth does not rely on diffusion or advection. Instead, the process of branching displaces the center of mass of the structure. Thus, it emerges as consequence of the finite extension and dynamics of the reactant (microtubules). The interplay between autocatalytic growth and fluxes driven by motors could lead to general principles of pattern formation and cytoskeletal organization in cells.

## Acknowledgements

We acknowledge A. Hyman, M. Loose, S. Grill, T. Quail, K. Ishihara, R. Farhadifar, G. Pigino, F. Jug, C. Norden and I. Patten for fruitful discussions and careful revision of the manuscript. We also thank J. Rosenberger for helping on the coding for laser ablation. We kindly thank R. Ohi and K. Ishihara for providing us the anti-MCAK antibodies and RanQ69L, respectively. This work was supported by the Human Frontiers Science Program (CDA00074/2014, to JB), EMBO (ALTF 483-2016, to DO), the ELBE postdoctoral program (BD), and a DIGS-BB fellowship provided by the DFG (FD).

